# Nearest-neighbor effects modulate *loxP* spacer DNA chemical shifts and guide oligonucleotide design for NMR studies

**DOI:** 10.1101/2021.08.13.456142

**Authors:** Nicole Wagner, Mark P. Foster

## Abstract

Cre recombinase catalyzes site-specific DNA recombination at pseudo-palindromic *loxP* sites through two rounds of strand cleavage, exchange, and religation. Cre is a potential gene editing tool of interest due its lack of requirements for external energy sources or host factors, as well as the fact that it does not generate potentially cytotoxic double-stranded DNA breaks. However, broader applications of Cre in editing noncanonical target sequences requires a deeper understanding of the DNA features that enable target site selection and efficient recombination. Although Cre recombines *loxP* DNA in a specific and ordered fashion, it makes few direct contacts to the *loxP* spacer, the region where recombination occurs. Furthermore, little is known about the structural and dynamic features of the *loxP* spacer that make it a suitable target for Cre. To enable NMR spectroscopic studies of the spacer, we have aimed to identify a fragment of the 34-bp *loxP* site that retains the structural features of the spacer while minimizing the spectral crowding and line-broadening seen in longer oligonucleotides. We report sequential backbone resonance assignments for *loxP* oligonucleotides of varying lengths and evaluate chemical shift differences, Δδ, between the oligos. Analysis of flanking sequence effects and mutations on spacer chemical shifts indicates that nearest-neighbor and next-nearest-neighbor effects dominate the chemical environment experienced by the spacer. We have identified a 16-bp oligonucleotide that adequately preserves the structural environment of the spacer, setting the stage for NMR-based structure determination and dynamics investigations.

## Introduction

Cre Recombinase is a site-specific tyrosine recombinase originating from bacteriophage P1. Its roles in the bacteriophage life cycle include circularization of the linear P1 genome after entry into the host cell and resolution of plasmid dimers^1^. Cre has become a widely used tool in genetic manipulation due to several advantageous features; it does not require external host proteins^2^ or energy sources^3^ to function, and unlike nucleases that are commonly applied in gene editing, it does not generate double-stranded DNA breaks^3^. Cre is therefore a promising tool for further applications in gene editing, including therapeutic applications toward resolving genetic mutations. However, its limited substrate scope is perhaps its most significant limitation. Thus far, efforts to develop Cre mutants to recognize new, therapeutically relevant target sites have been conducted through many rounds of directed evolution^4,5^, and the resulting recombinases have still shown off-target activity on human genomic sequences in some studies^6^. Rational design of new mutants could be a more efficient approach. However, in order to engineer Cre mutants to recognize new target sites with high specificity, it is first necessary to deepen our understanding of how Cre selects and recognizes its substrate sequence.

The canonical target site for Cre is called *loxP*. The *loxP* site consists of two 13 bp inverted repeat sequences called Recombinase Binding Elements (RBEs) flanking an 8 bp asymmetric spacer^7^. One Cre protomer binds at each RBE, and then two such Cre_2_-*loxP* complexes come together to form the tetrameric synaptic complex (Fig. 1A). Cre then proceeds to recombine the DNA through two rounds of strand cleavage, exchange, and religation^3^. Recombination occurs at the spacer, with the central 6 bp forming the crossover region. Homology between the two *loxP* sites in the tetramer is required for efficient recombination^7^. Furthermore, strand exchange occurs in an ordered fashion, where the so-called “bottom strands” of the two *loxP* sites are preferentially cleaved and exchanged first, followed by the “top strands” in the second round^8^.

**Figure 1.**
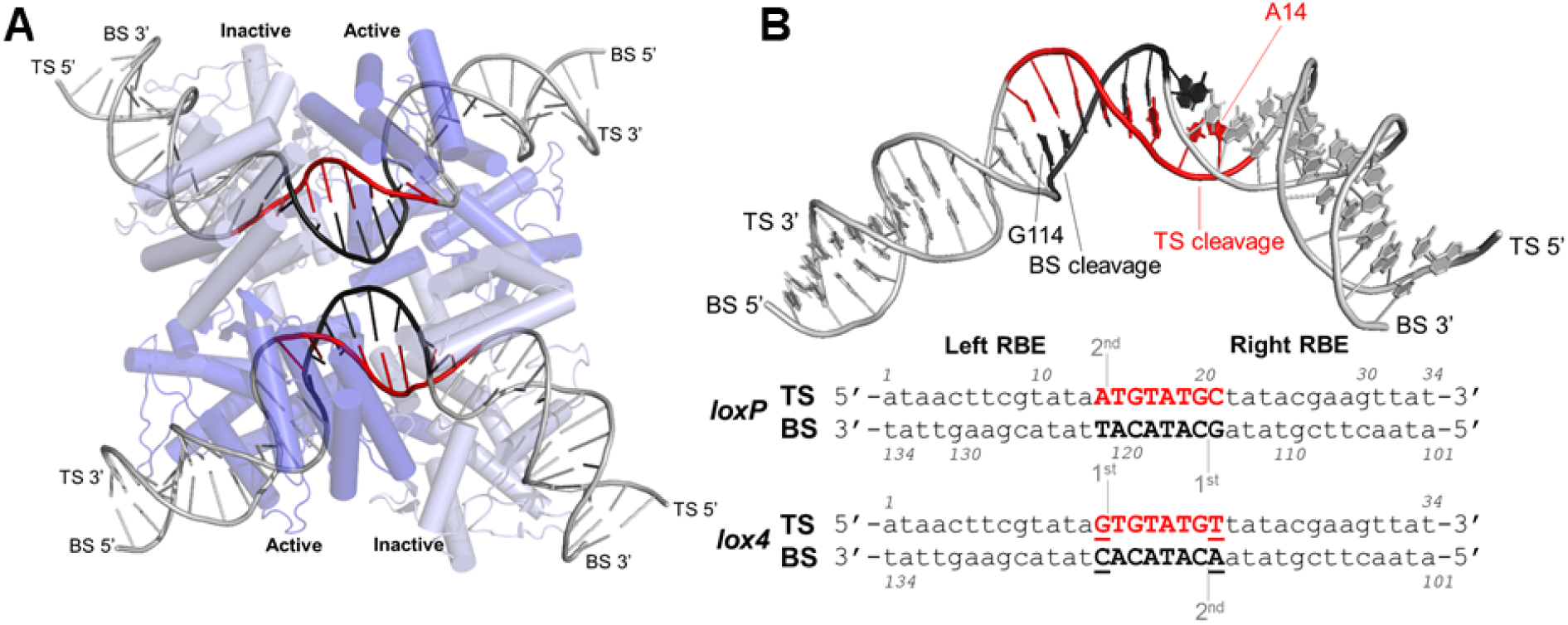
Cre and *loxP*. A) Synaptic complexes (PDB:2HOI) of four Cre (K210A) protomers, two of which are active at a time, assembled on two *loxP* sites, and featuring a bend in the spacer DNA^11^. In this complex, the protomers on the “bottom strands” (BS; black) are “Active” and poised for cleavage; the protomers on the “top strands” (TS; red) are “Inactive”. B) Top: *loxP* DNA duplex from PDB: 2HOI. Bottom: sequences of *loxP*, the native substrate for Cre, and of *lox4*. Mutated positions in *lox4* are underlined. Strand cleavage on both duplexes takes place after the 14th nucleotide of each *lox* site. For *loxP*, the reaction proceeds preferentially through bottom strand cleavage and exchange, followed by the top strand. *lox4* shows a reversed order of strand cleavage and exchange^8^. Cleavage sites and their order are indicated.

Cre can recombine *lox* sites with noncanonical spacers (as long as spacer homology is maintained between the pair of target sites), but none with the same efficiency as the WT spacer^9^. A spacer variant called *lox4* has been engineered to interconvert the outermost base pairs of the spacer at each end (the positions 5’ to the scissile phosphates that are cleaved). It has been shown that this interconversion causes *lox4* to be recombined with a reversed order of strand exchange (Fig. 1B), albeit with twofold lower efficiency than *loxP*^8^. Despite the demonstrated impacts of spacer mutations, Cre makes few direct base-specific contacts with the spacer^10^. This may implicate intrinsic flexibility of the spacer in promoting site-specific recognition and efficient recombination by Cre, and in directing the order of strand exchange. A crystal structure of the pre-cleavage tetrameric synaptic complex shows that a kink at one end of the spacer coincides with activation of the other end for cleavage^11^, while mobility shift assays suggest asymmetric bending of the spacer in pre-synaptic complexes^12^. These underscore the likely role of the asymmetric spacer sequence in directing the asymmetry of the synaptic complex (in which one protomer at each *loxP* site is active at a time) and the order of strand exchange^13^.

Currently, most detailed structural knowledge of Cre and *loxP* comes from static crystal structures. However, nanosecond-timescale MD simulations have suggested that the base-pair steps flanking the bottom strand (BS) cleavage site of the WT spacer are more flexible than those flanking the top strand (TS) cleavage site, resulting in opening of the minor groove and positioning of the cleaving protomer. Conversely, the steps flanking the TS cleavage site in *lox4* were shown to be slightly more flexible than those flanking the BS cleavage site, which may help explain the reversed strand exchange order^14^. The data from the MD simulations support the notion that intrinsic flexibility within and around the *loxP* spacer plays key roles in recognition and recombination by Cre. The existing knowledge gained from MD simulations can be expanded upon by performing NMR measurements of spacer DNA structure and dynamics.

Here, we describe work to identify a *loxP* spacer oligo that minimizes resonance overlap by truncating portions of the flanking regions while still maintaining a chemical environment that resembles the environment the spacer experiences in longer oligos. NMR chemical shifts report on the unique environment experienced by a particular nucleus. Therefore, we collected chemical shift data for the *loxP* spacer with varying compositions of flanking regions and degrees of native sequence context. We report the effects of flanking sequences on the chemical shifts of base H6/H8 and sugar H1’ nuclei within the spacer. We also report the extent of chemical shift differences, Δδ, for all residues in the *lox4* spacer compared to *loxP*. Our findings show that alterations in the flanking regions, as well as the mutations in the outermost base pairs in *lox4*, introduce primarily local structural perturbations. We also compare our observed chemical shifts with predicted chemical shifts in order to identify nuclei within the *loxP* spacer that may experience structural context differing from B-form DNA.

Our work demonstrates the use of a “divide and conquer” approach^15^ to study a portion of the 34 bp *loxP* site. Nucleic acids experience more resonance overlap than proteins, as there are only 4 different canonical residue types in DNA or RNA compared to 20 in proteins. The redundancy of DNA chemical shifts can necessitate either site-specific isotope labeling at the positions of interest, or development of smaller oligos that preserve the regions of interest while removing extraneous regions. However, it is important to verify that truncation of flanking regions does not significantly alter the chemical environment experienced by the region of interest. Our findings therefore have more general implications for the design of NMR constructs representing regions of interest within larger DNAs.

## Methods

### DNA oligonucleotide sample preparation

Single-stranded DNA oligonucleotides were purchased from Integrated DNA Technologies (Coralville, IA) with standard desalting purification. The lyophilized oligos were resuspended in MilliQ water. Equimolar quantities of the complementary strands were combined, and concentrated stocks of Tris (pH 7) and NaCl were added to concentrations of 10 mM and 100 mM, respectively. Annealing was performed by placing the mixture in a water bath at 95 °C, shutting off the heat, and cooling gradually in the water bath overnight. The annealed oligos were further purified using anion exchange chromatography with a HiTrap QHP column (Cytiva). FPLC fractions were concentrated using a Macrosep 1k MWCO centrifugal filter (Pall) at 4 °C.

### NMR experiments

Samples were exchanged into NMR buffer (10 mM D11-Tris, 100 mM NaCl, pH 7) using an Amicon 0.5 mL 3k MWCO centrifugal filter (Millipore). Sample concentrations ranged from ~250 μM to ~1.4 mM at 500 μL volume. D_2_O was added to 10% (v/v), DSS was added as an internal reference to a concentration of 50 μM, and 0.02% NaN_3_ was added to prevent microbial growth. NMR data were recorded at 25 °C using a Bruker Avance III HD Ultrashield 600 MHz spectrometer equipped with a 5 mm triple-resonance inverse (TXI) cryoprobe with z-gradients. Data were processed and visualized with NMRFx^16^, NMRViewJ^17^, and Topspin (Bruker). Typical 2D NOESY spectra were collected with 64 scans, spectral widths of 13227.51 x 13192.61 Hz, and a mixing time of 200 ms. NOESY spectra were apodized with a 90° shifted squared sine bell in F2 and a 54° shifted squared sine bell in F1, and data were zero-filled to 8192 x 2048 points. Proton chemical shifts were assigned from 2D NOESY spectra using standard sequential walk approaches^18^. 2D TOCSY spectra were collected with 64 scans, spectral widths of 14423.08 x 14409.22 Hz, and a mixing time of 75 ms. TOCSY spectra were apodized and zero-filled in the same manner as the NOESY spectra, and were used to verify cytosine H5-H6 correlations.

### T_m_ prediction

Oligo melting temperatures, T_m_, were predicted using OligoCalc^19^, with adjustments for salt concentration and nearest neighbor effects, using the following equation, wherein values of ΔH and ΔS are estimated from nearest-neighbor parameters of the input sequence^20^. The values used for [oligo] and [Na^+^] were 25 μM and 100 mM, respectively (the conditions of our thermal melt experiments).

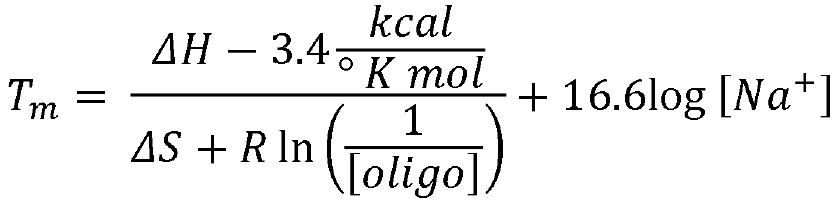

### UV-Vis thermal melts

Samples were diluted in NMR buffer (10 mM D11-Tris, 100 mM NaCl, pH 7) to a concentration of 25 μM and placed into a 1 mm path length quartz cuvette (Starna Cells). NMR buffer with no DNA was used as a blank. Melting curves were acquired by measuring absorbance at 260 nm over a range of temperatures using a Hewlett-Packard 8452A Diode Array Spectrophotometer with UV-Visible ChemStation software. Temperature ramp was performed in 1 °C increments from 15 °C to 70 °C (10- and 12-mer) or 80 °C (16- and 22-mer) with a hold time of 30 seconds at each increment. Melting temperatures were determined by fitting to a two-state model^21,22^.

### 3D coordinates and Δδ mapping

3D models of the *loxP* spacer oligonucleotides as B-form DNA were generated using the 3D-DART web server^23^, and hydrogen atoms were added to the structure using MolProbity^24,25^. Δδ values for the H6/H8 and H1’ protons in shorter oligos relative to the 22-mer were mapped onto the spacer structure as a linear gradient from white (0.010 ppm) to red (0.402 ppm). Δδ values for the H6/H8 and H1’ protons in the *lox4* 16-mer relative to the *loxP* 16-mer were mapped onto the structure as a linear gradient from white (0.000 ppm) to red (0.111 ppm).

### Chemical shift predictions

Predicted chemical shifts for double-helical B-form DNA and random coil DNA were calculated from DSHIFT^26^. The Altona method was used within DSHIFT for prediction of double-helical B-form chemical shifts^27^.

## Results

### Sequential assignments of *loxP* oligos

In order to minimize complications from overlap of NMR resonances and effects of anisotropic tumbling on relaxation rates, we set out to determine the smallest fragment of the *loxP* sequence that would retain the structural and dynamic features of the spacer in longer DNA oligonucleotides. Because they are generally the best resolved resonances in DNA, we used the purine H8, pyrimidine H6 and sugar H1’ protons as the reporters of local environment. We thus assembled a series of DNA oligonucleotides comprising the 8 bp spacer, 5’-d(ATGTATGC)-3’, plus flanking sequences of varying length (for a total of 10, 12, 16, or 22 base pairs) and composition (Fig. 2).; flanking sequences were chosen to limit end-fraying by inclusion of GC pairs, to maximize native sequence context, or both.

**Figure 2.**
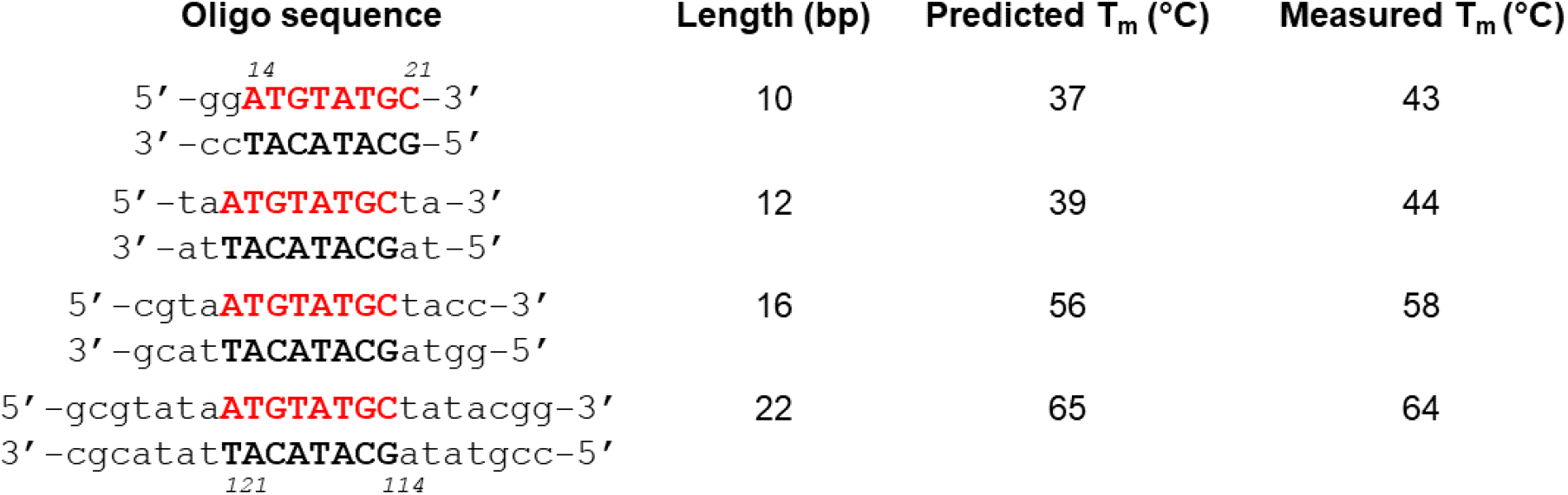
Sequences of the 10-mer, 12-mer, 16-mer, and 22-mer *loxP* constructs used in this paper. Theoretical T_m_ values were predicted using OligoCalc^19^ and experimental T_m_ values were determined from UV-Vis thermal melts (see Methods).

#### 10-mer

Because its smaller size resulted in narrow NMR resonance lines and reduced overlap, leading to simple spectra, we first made resonance assignments of the 10-mer oligo. This oligo consisted of the 8 bp spacer, with a 2 bp GC clamp added at the AT-rich end of the spacer to mitigate end fraying (Fig. 2). The H6/8-H1’/Cyt H5 region of the NOESY spectrum showed narrow lines and generally well-dispersed signals (Fig. S2). We observed all expected intra- and inter-residue H6/8-H1’ NOEs for the spacer region of the oligo (a total of 31 NOEs within the spacer), as well as most of the expected NOEs for the flanking region. An inter-residue H8 to H1’ NOE between G13 and G12 of the GC clamp was not plainly visible due to presumed overlap in their H1’ chemical shifts. Other resonances that were used to support the sequential assignments for the 10-mer (and the longer oligos) were NOEs from thymine methyl (H7) protons and cytosine H5 protons to their own H6 and the H6/8 of the preceding (5’) nucleotide. Overall, we obtained complete H6/8 and H1’ assignments for the entire spacer for this 10-mer oligonucleotide.

#### 12-mer

The 12-mer oligo consisted of the 8 bp spacer flanked by 2 bp of native sequence context on each side, with no GC clamps (Fig. 2). The H6/8-H1’/Cyt H5 region of the NOESY displayed sharp lines (Fig. S3), although the spectrum was not as well-dispersed as that of the 10-mer. We expect 32 NOEs within the 8-bp spacer (two strands each featuring intra- and inter-nucleotide NOEs), but only 26 were apparent before taking into consideration resonance overlap (particularly within the adenine H8 and thymine H6 regions). Compared to the 10-mer, the signals for the 12-mer were somewhat less dispersed, likely due to the palindromic nature of the flanking regions – even some signals that were well-separated in the NOESY of the 10-mer (such as T15/T19) displayed overlap for the 12-mer. Nevertheless, we were able to assign chemical shifts for all H6/8 and H1’ protons within the spacer using the sequential walk approach (Fig. S3). The flanking regions were only partially assigned, as resonance overlap made it difficult to obtain full unambiguous assignments for the flanking regions.

#### 16-mer

The 16-mer oligo contained the same native sequence context as the 12-mer (2 bp flanking each end of the spacer) with an additional 2 bp GC clamp introduced at each end (Fig. 2). The H6/8-H1’/Cyt H5 region of the NOESY spectrum was characterized by relatively narrow lines and well-dispersed resonances (Fig. 3, Fig. S1). Some spectral crowding was present among cytosine and thymine H6 protons, but the degree of crowding was not severe enough to be unresolvable by standard sequential walk approaches. The palindromic flanking regions did not cause as severe of resonance overlap as in the 12-mer, because the GC clamps were asymmetric, resulting in different chemical shifts for the 5’ and 3’ flanking residues. Complete H6/8 and H1’ resonance assignments could be obtained for the spacer (Fig. 3, Fig. S1) and for the flanking regions.

**Figure 3.**
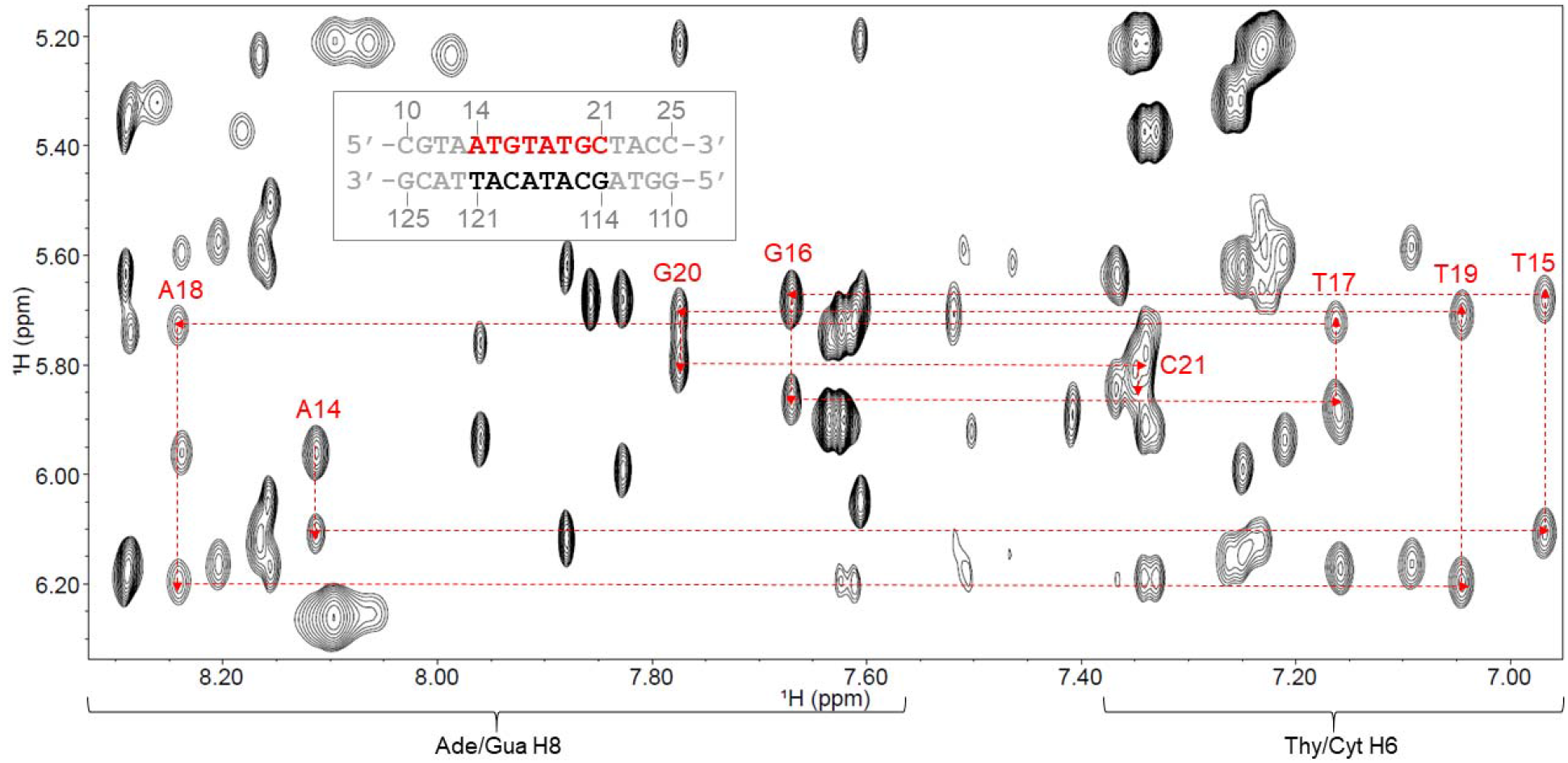
Aromatic H6/H8 to anomeric H1’ “NOESY-walk”^18^ for the top strand (TS) of the 16-mer construct. Labels indicate the residue to which the H6 or H8 belongs. The numbering scheme is shown in the inset.

#### 22-mer

The 22-mer oligo contained the spacer flanked by 6 bp of native sequence context on each side, as well as one additional GC base pair on each end to mitigate end fraying. The length of the oligo combined with the palindromic nature of the flanking regions resulted in significant resonance overlap, which made it challenging to assign the spacer *de novo* from the 22-mer NOESY spectrum. Furthermore, broad lines were observed for many resonances, compared to the narrow lines seen for the shorter oligos. The top strand was more easily assigned than the bottom strand, as it had fewer resonances located in regions of severe overlap. Severe resonance overlap rendered tentative our assignments for the bottom strand, particularly for C115 (H6 and H1’), A116 (H8 and H1’), T117 (H6), A118 (H1’), C119 (H6 and H1’), A120 (H6 and H1’), and T121 (H6). It was possible to infer chemical shift assignments for all H6/8 and H1’ protons within the spacer (Fig. S4), but the palindromic flanking regions were left unassigned.

### Flanking sequences have the largest effect on spacer chemical shifts in the outer positions of the spacer

With complete spacer H6/8 and H1’ assignments for all oligos, we compared the chemical shifts in order to determine how much of the flanking sequences should be retained in order to accurately preserve the chemical environment experienced by the spacer, using the largest oligo as reference (the 22-mer). To assess how the chemical environment of the spacer was impacted by shortening the flanking sequences, we compared the H6/8 and H1’ chemical shifts of the shorter oligos to the 22-mer (Fig. 4) and calculated the Δδ, defined as the difference in ^1^H chemical shift from the 22-mer. As expected from nearest-neighbor and next-nearest-neighbor effects^27,28^, compared to the 22-mer, the largest Δδ values were observed closer to the ends of each spacer (primarily in the first two base pairs at either end), and decreased towards the center of the spacer (Fig. 5).

**Figure 4.**
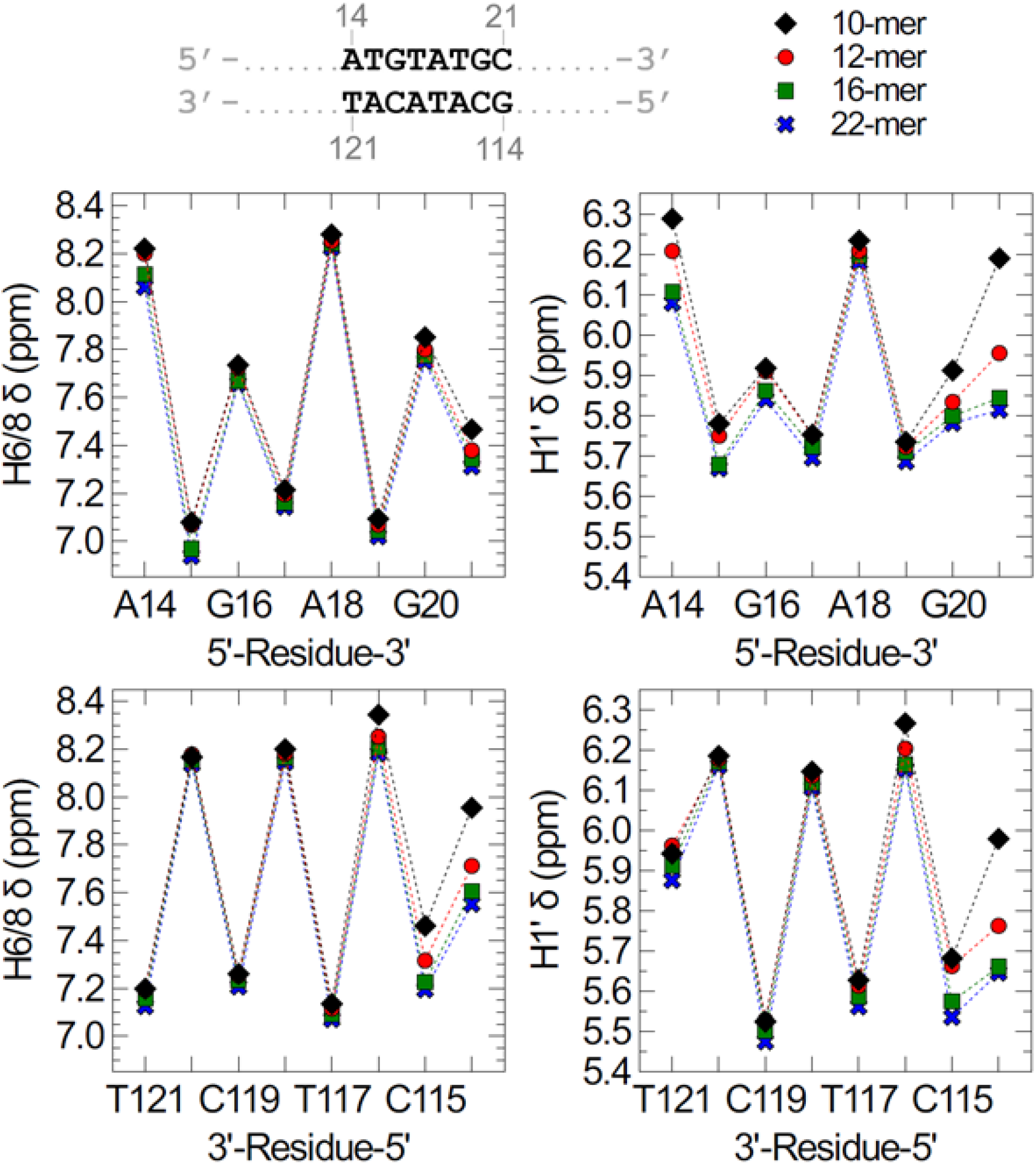
H6/H8 (left) and H1’ (right) chemical shifts of *loxP* spacer nucleotides. Top, TS chemical shifts according to sequence in a 5’-3’ direction; bottom, BS chemical shifts are plotted in a 3’ to 5’ direction such that base-paired partners are directly across from one another. Black = 10-mer; red = 12-mer, green = 16-mer; blue = 22-mer.

**Figure 5.**
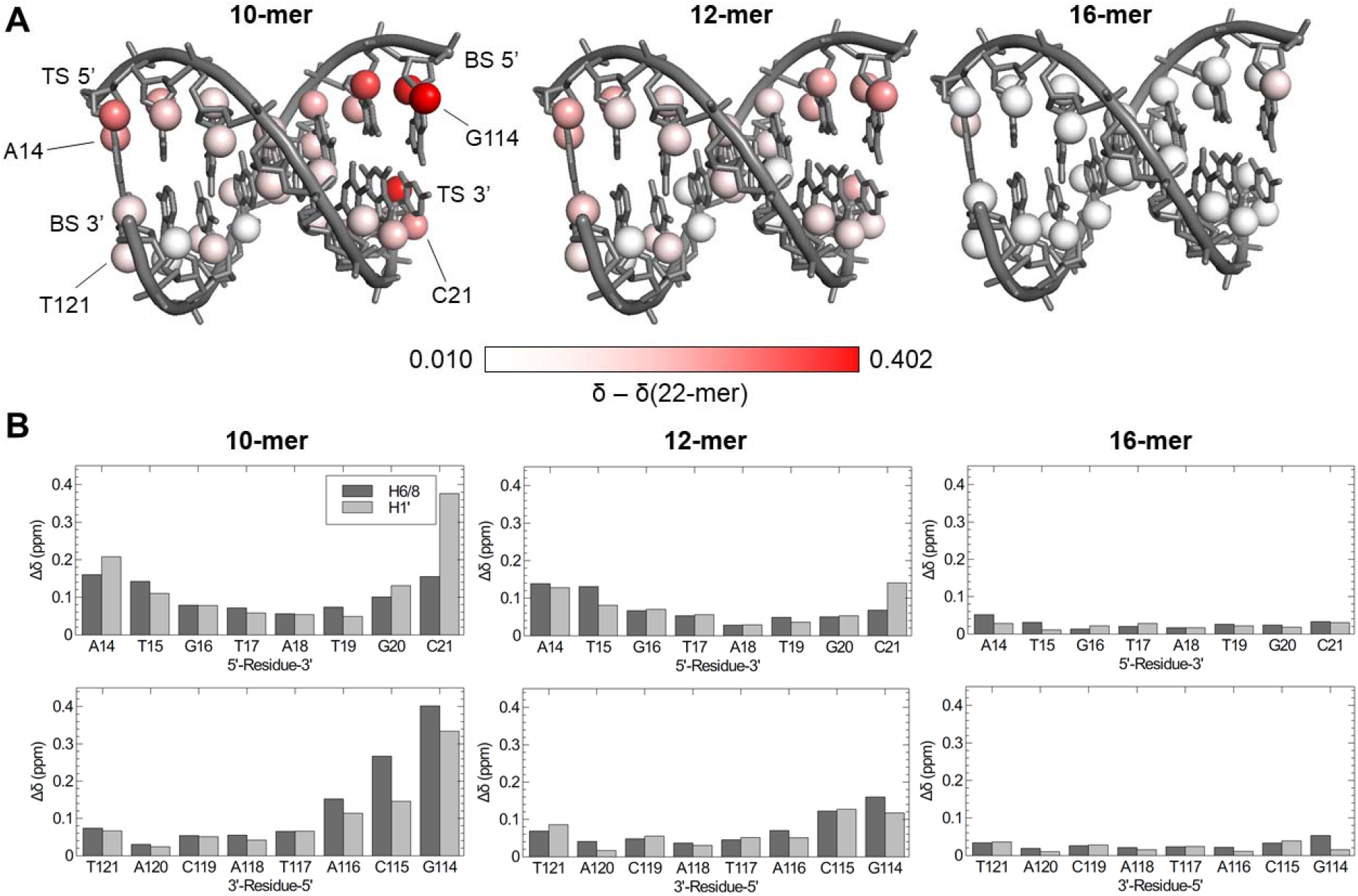
Chemical shift differences (Δδ values) of spacer oligonucleotides, relative to the 22-mer. A) Δδ mapped to a 3D model of the *loxP* spacer. H6/8 and H1’ atoms are shown as spheres and colored using a linear white-to-red gradient, with red indicating the largest value. In order from left to right: Δδ values of the 10-mer, 12-mer, and 16-mer oligos. B) Δδ values relative to 22-mer chemical shifts shown as bar graphs. In order from left to right: values for the 10-mer, 12-mer, and 16-mer oligos. TS values are shown in the top plot; BS values are plotted below such that base-paired partners are directly across from one another. For comparative purposes, the y-axes are the same for each plot.

#### 10-mer

In the 10-mer, the C21 and G114 H1’ and H6/8 resonances (at the end of the spacer) in particular are shifted significantly downfield compared to their location in any other oligo we studied, with Δδ as large as ~0.4 ppm. Δδ values decreased to ~0.27 and then ~0.15 ppm at the 2^nd^ and 3^rd^ base pairs in from the end (Fig. 4). Most resonances in the 10-mer were shifted more downfield than the same resonances in the other oligos. This may be attributable to an increased shift toward a single-stranded (random coil) population^29^, consistent with its lower melting point (Fig. 2). The RMSD of the Δδ’s between the 10-mer and 22-mer was 0.153 for H6/8 protons, 0.156 for H1’ protons, and 0.154 for H6/8 and H1’ protons combined.

#### 12-mer

As with the 10-mer, all of the largest Δδ values were located at the ends of the spacer, and generally decreased moving closer to the center (Fig. 5). Although the overall pattern was unchanged from the 10-mer, the Δδ values at the ends of the spacer were smaller in magnitude than for the 10-mer, owing to the native sequence context that was introduced in the 12-mer. The 12-mer resonances still trend toward a downfield chemical shift compared to the 16-mer and 22-mer. As with the 10-mer, this could be attributed to an increased shift toward a small single-stranded population compared to the longer oligos due to a lower melting point (Fig. 2). The RMSD of the Δδ’s between the 12-mer and 22-mer was 0.0836 for H6/8 protons, 0.0801 for H1’ protons, and 0.0819 for H6/8 and H1’ protons combined.

#### 16-mer

The addition of GC clamps at each end of the 16-mer helped to reduce the Δδ values at the ends of the spacer, compared to the Δδ’s observed for the same resonances in the 12-mer (Fig. 5). As the non-native GC clamps are 3 bp away from the spacer, they should not exert nearest-neighbor or next-nearest-neighbor effects on the chemical environment experienced by the spacer, but mitigation of end fraying may have introduced a more native-like structure than in the 12-mer (see Discussion). Overall, the 16-mer showed the smallest Δδ values from the 22-mer for each of the positions we studied. As with the other oligos, the largest Δδ values were observed at the ends of the spacer; however, even these Δδ values were smaller in magnitude than in the other oligos, with all Δδ values smaller than 0.1 ppm relative to the 22-mer, and with most being smaller than 0.05 ppm (Fig. 5B). The RMSD of the Δδ’s between the 16-mer and 22-mer was 0.0301 for H6/8 protons, 0.0237 for H1’ protons, and 0.0271 for H6/8 and H1’ protons combined.

### Comparisons with B-form and random coil chemical shift predictions

We compared experimentally determined H6/H8 and H1’ chemical shifts for each oligonucleotide to B-form and random coil (single-stranded) shifts predicted from empirical nearest-neighbor effects^26,27,29^ (Fig. S7). For the 12-, 16-, and 22-mer oligos, the H6/8 proton with the largest deviation from B-form predictions was at position G16, and the second-largest was at A120. In the 10-mer, G16 and A120 were tied as the largest H6/8 deviation from B-form predictions. In all oligos, G16 H8 and A120 H8 were upfield from the predicted B-form values (shifted away from random coil). Furthermore, the difference between observed and predicted values at these positions often increased with the length of the oligo (to an observed value further from random coil). The H1’ protons with the largest deviations from the predictions were at positions T17, C115, and C119 in the 10-mer; C115 and C119 in the 12-mer; C21, G16, A116, and C119 in the 16-mer; and G16, C21, and A116 in the 22-mer. Overall, cytosine H1’ protons were overrepresented in this group, and they were often downfield of the predicted B-form shifts (towards random coil). However, in the 16-mer and 22-mer, C21 showed a large upfield shift from the predicted B-form value. T17 H1’ in the 10-mer was also downfield of the predicted value, while G16 H1’ and A116 H1’ in the 16-mer and 22-mer were upfield.

Although many of the predictions for B-form DNA were fairly accurate, the RMSDs between observed and predicted values often increased with oligo length. The RMSDs between observed and predicted B-form values for all *loxP* oligos are summarized in Table 1. Comparisons with random coil predictions showed that the 10-mer and 12-mer were often shifted towards random coil more so than the 16-mer or 22-mer. The RMSDs between observed and predicted random coil values for all *loxP* oligos are summarized in Table 2. Although nearly all of our observed chemical shifts were upfield of the random coil predictions (with the single exception of A116 H8 in the 10-mer), the differences between random coil and observed chemical shifts were often smaller in the shorter oligos. A rare exception is the T121 H1’, where the 16-mer chemical shift is slightly closer to the random coil prediction than in the 10-mer. As discussed above and as shown in Fig. 2, the lower T_m_ values for the 10-mer and 12-mer may cause their chemical shifts to be weighted towards a larger random coil population than the longer oligos.

**Table 1.**
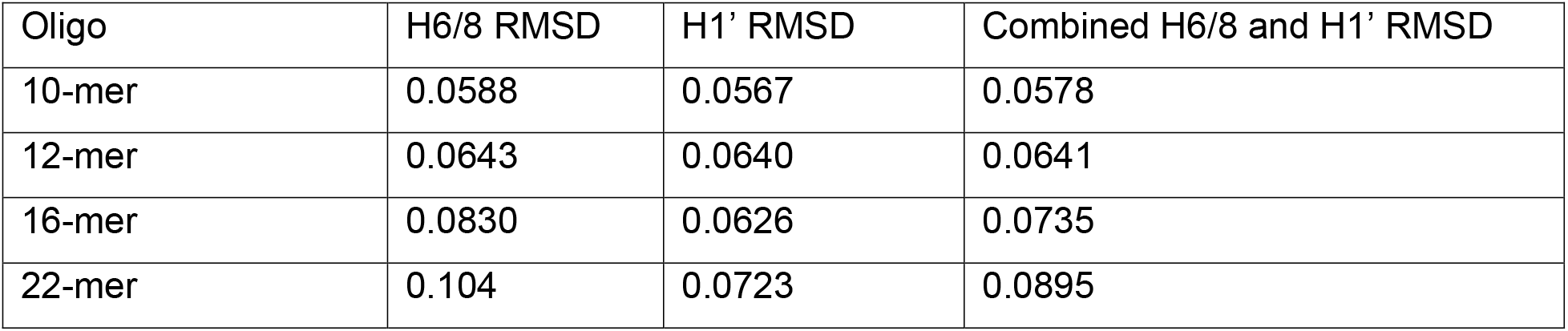
RMSDs between observed chemical shifts and predicted B-form duplex chemical shifts for all *loxP* oligos.

**Table 2.** RMSDs between observed chemical shifts and predicted random coil chemical shifts for all *loxP* oligos.

| Oligo | H6/8 RMSD | H1' RMSD | Combined H6/8 and H1' RMSD |
| --- | --- | --- | --- |
| 10-mer | 0.147 | 0.263 | 0.206 |
| 12-mer | 0.177 | 0.262 | 0.224 |
| 16-mer | 0.220 | 0.307 | 0.267 |
| 22-mer | 0.247 | 0.330 | 0.291 |

**Table 3.** RMSD values of *lox4* chemical shifts from *loxP* at nearest-neighbors (NN) of the mutated sites, next-nearest-neighbors (NNN) of the mutated sites, and non-NN or NNN positions.

| Position Type | H6/8 RMSD | H1' RMSD | Combined H6/8 and H1' RMSD |
| --- | --- | --- | --- |
| NN | 0.0654 | 0.0945 | 0.0812 |
| NNN | 0.0615 | 0.0395 | 0.0517 |
| Non-NN or NNN | 0.0244 | 0.0111 | 0.0190 |

### lox4 mutations have the largest effect on spacer chemical shifts ≤ 2 bp away from mutated residues

To understand how spacer sequence elements can alter recombination, we obtained resonance assignments for a 16-mer oligonucleotide bearing the *lox4* spacer sequence, in which the outermost GC and AT base pairs at each end of the spacer are swapped (Fig. 1B). As with the *loxP* 16-mer, assignments for all H6/8 and H1’ protons could be obtained for the spacer (Fig. S5) and the flanking regions.

Comparison of the H6/8 and H1’ Δδ values for the *lox4* 16-mer relative to the equivalent positions in the *loxP* 16-mer showed that, within the spacer, significant chemical shift changes were limited to ~2 bp from the site of mutation (Fig. 6). Ignoring the sites of mutations, the largest Δδ values were generally localized up to 2 bp away from the mutated positions. Spacer positions that were neither nearest-neighbors (NN) nor next-nearest-neighbors (NNN) to the mutated sites showed considerably smaller RMSDs from *loxP* compared to NN and NNN positions. These data suggest that the mutations in the *lox4* spacer introduce primarily local structural changes that do not propagate dramatically throughout the entire spacer.

**Figure 6.**
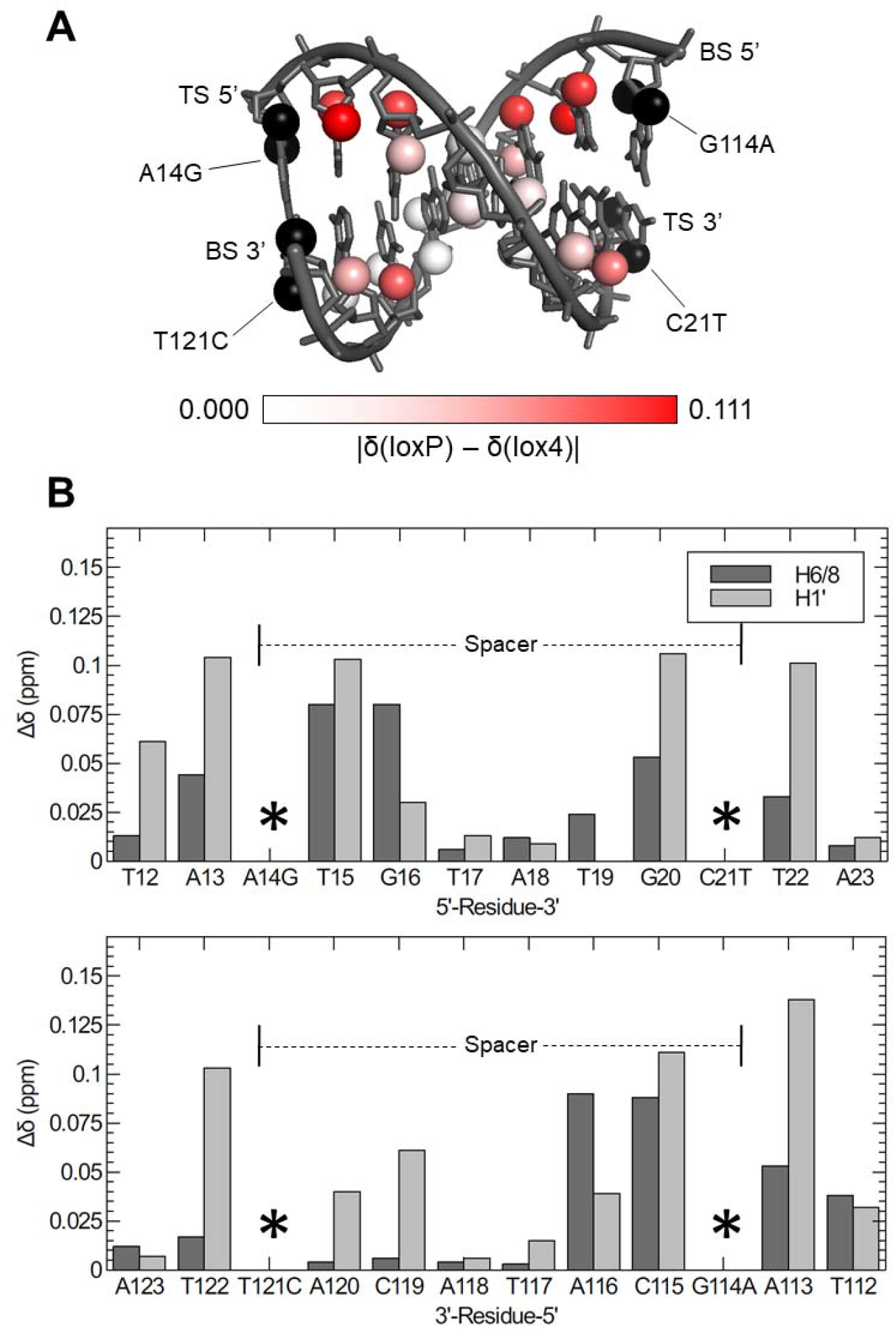
Effects of the *lox4* mutations on spacer and flanking sequence H6/H8 and H1’ chemical shifts. A) Δδ values for the spacer of the *lox4* 16-mer relative to the *loxP* 16-mer, mapped to a 3D model of the WT *loxP* spacer. H6/8 and H1’ atoms are shown as spheres and colored using a linear white-to-red gradient, with red indicating the largest Δδ. B) Δδ values of the *lox4* spacer relative to *loxP*, shown as bar graphs. Asterisks denote mutated positions. TS values are shown in the top plot; BS values are plotted below such that base-paired partners are directly across from one another. Also shown are Δδ values of the native sequence regions flanking the spacer in *lox4* relative to *loxP*.

In the native sequence positions of the flanking regions, the H1’ chemical shifts often appeared more sensitive to the mutations than H6/8 chemical shifts (Fig. 6B). With the exception of A123 and T112, each H1’ Δδ value was > the H6/8 Δδ for the same residue. In contrast, our other Δδ measurements (*lox4* spacer vs. *loxP* spacer, or shorter *loxP* oligos vs. 22-mer) did not show a strong consistent pattern in whether the H6/8 or H1’ shifts were more sensitive to environmental changes. As expected based on nearest-neighbor effects, the native sequence positions immediately adjacent to the spacer showed larger Δδ values than the positions two base pairs away from the spacer.

## Discussion

The goal of this study is to identify a tractable DNA oligonucleotide sequence for NMR-based structural and dynamic analysis of the asymmetric *loxP* spacer, which directs the order of preferential strand exchange by Cre recombinase, and of a *lox4* variant, which features reversed order of strand exchange. Backbone H6/H8 and H1’ chemical shifts were assigned and compared as proxies for DNA structure and dynamics for oligos comprising the 8-bp spacer plus additional flanking sequences, with total lengths of 10-, 12-, 16-, and 22-bp (Figure 2). As expected from nearest-neighbor effects on DNA structure and stability^27,28^, effects of alterations in the flanking sequences were primarily limited to the next-nearest neighbors and did not propagate throughout the spacer.

In the 22-mer, the 7-bp regions on each side of the spacer were almost entirely palindromic, except for the terminal GC base pair added to each end to mitigate end fraying. The palindromic nature of these regions led to significant resonance overlap, and as a result, the flanking regions were left unassigned in the 22-mer. Furthermore, resonance overlap in the 22-mer was severe enough to render tentative some of the spacer assignments, as chemical shifts of some spacer residues also occurred within overlapped regions (see Results). The lower precision of the 22-mer assignments presents a limitation in our comparisons of the shorter oligos with the 22-mer chemical shifts, as Δδ values were calculated relative to the 22-mer.

The 16-mer contained the same native sequence region as the 12-mer, with an asymmetric 2-bp GC clamp introduced at each end. Interestingly, the addition of GC clamps, despite being non-native, brought the chemical shifts at the ends of the spacer into better agreement with the 22-mer. This suggests that the AT-rich flanking sequences of the 12-mer may have resulted in significant end-fraying. End-fraying in the flanking regions of the 12-mer represents a deviation from the native structure despite possessing the native sequence, and therefore impacts the chemical environment experienced by the outer regions of the spacer.

Analysis of the differences between observed chemical shifts and B-form and random coil predictions highlighted unique features near the cleavage sites. In each of the oligos, the G16 H8 proton in a 5’-TGT-3’ context showed the maximum deviation from predicted double-helical B-form chemical shifts (Figure S7). In the 12-, 16-, and 22-mer, G16 H8 had the largest deviation of any H6/8 proton (0.14, 0.19, and 0.20 ppm, respectively); in the 10-mer, only A120 H8 had a deviation of equal magnitude (0.12 ppm). This may reflect limitations of the training sets used for chemical shift predictions^27^; however, in the crystal structure of the tetrameric synaptic complex with cleavage-deficient Cre-K201A, G16 is located adjacent to the kink in the spacer^11^. Additionally, the T15pG16 step was identified as a flexible step in 15-ns MD simulations^14^. We can therefore speculate that G16 H8 may be able to experience conformations that differ from idealized B-form DNA, such as altered helicoidal parameters or groove widths. G16 was also among the positions with the largest differences between observed and predicted H1’ chemical shifts in the 16-mer and 22-mer, lending further support to the structural relevance of G16. Furthermore, the largest H6/8 and H1’ deviations from duplex predictions in *lox4* occurred at position A116 (0.18 and 0.13 ppm, respectively; Fig. S8), which would be the equivalent to position G16 in a top-strand cleavage scenario, assuming the opposite end of the spacer adopts a bend equivalent to that seen in the pre-cleavage synaptic complex when poised for bottom-strand cleavage^11^.

The A120 H8 represented the second-largest H6/8 deviation from predicted values in the 12-, 16-, and 22-mer. 15-ns MD simulations also identified the C119pA120 step as a flexible step^14^. In bottom-strand cleavage scenarios (the first cleavage step in the mechanism^8^), the “RK motif” of Cre (residues R118, R121, and K122) clamps around the region of the spacer containing A120 in order to narrow the minor groove, thus discouraging cleavage at this end of the spacer and promoting cleavage at the opposite end^14^. Therefore, A120 H8 may also experience alternate conformations that differ from idealized B-form DNA; for example, this region of the spacer may be predisposed toward a narrowed minor groove. The equivalent position to A120 in a top-strand cleavage scenario is G20, which also experienced reasonably large deviations from its predicted H8 shift in the 16-mer and 22-mer. In *lox4*, the size of the H6/8 deviations from predicted duplex values at G20 and A120 were nearly inverted compared to *loxP* (Fig. S8). In *loxP*, the G20 and A120 deviations were 0.10 and 0.13, respectively; in *lox4*, they were 0.14 and 0.11, respectively. This is an interesting observation, considering the reversed order of strand cleavage in *lox4*.

The RMSDs between observed chemical shifts and predicted double-helical B-form chemical shifts generally increased with the length of the oligo. This suggests that the addition of native flanking sequence context was able to induce structural features in the spacer that set it apart from idealized B-form DNA. Although the 10-mer oligo had the simplest spectra with minimal resonance overlap, the absence of native sequence context would therefore make it a poor representation of the structural and dynamic environment experienced by the spacer.

Finally, we evaluated the impact of the spacer mutations in *lox4* on the non-mutated positions. The largest spacer Δδ’s induced by the mutations were at the nearest-neighbors and next-nearest-neighbors of the mutated positions. RMSD values between *loxP* and *lox4* were larger at NN and NNN positions, and smaller for the spacer positions that were neither NN nor NNN to the mutated positions. This principle appears consistent with our study of Δδ’s in the shorter *loxP* oligos relative to the 22-mer, where we observed that positions distant from flanking sequence modifications were less dramatically impacted than the outer regions of the spacer. Furthermore, these observations are consistent with the expected dominant effect of nearest-neighbor and next-nearest-neighbor interactions in defining DNA strucure^27,28^.

## Conclusions

Cre Recombinase possesses unmet potential as a gene editing tool due to its lack of requirements for external energy sources or host factors^2^, as well as the fact that its recombination products do not require repair of double-stranded DNA breaks^3^. However, broader applications of Cre in editing noncanonical target sequences requires a deeper understanding of the *loxP* DNA features that enable target site selection and efficient recombination. The *loxP* spacer is of particular interest, as it is the site of recombination and because Cre makes few base-specific contacts with it. We sought to identify a minimal DNA fragment that could maintain a native-like environment for the *loxP* spacer while reducing the spectral crowding and line-broadening effects observed in NMR spectra of longer DNAs. Based on our findings, it appears that the 16-mer oligo achieves a balance between native sequence context and well-resolved spectra. Our oligo design and chemical shift assignments set the stage for future structural and dynamic measurements of the *loxP* and *lox4* spacers, which we expect to shed light on the basis for preferential order of strand cleavage by Cre Recombinase.

## Accession Codes

NMR chemical shift data have been deposited in the Biological Magnetic Resonance Data Bank (accession nos.: *loxP* 10-mer, 51032; *loxP* 12-mer, 51035; *loxP* 16-mer, 51036; *loxP* 22-mer, 51037; *lox4* 16-mer, 51047).

## Acknowledgements

We thank Chunhua Yuan and Alex Hansen of The Ohio State University Campus Chemical Instrument Center for assistance with NMR experimental setup, Kye Stachowski for providing the 22-mer oligo design, and the Foster lab members for helpful discussions and critical reading of the manuscript. This study was supported by NIH grant R01 GM122432 (to M.P.F.). N.W. was supported in part by a fellowship from The Ohio State University Graduate School.

## Supporting Information

**Figure S1.**
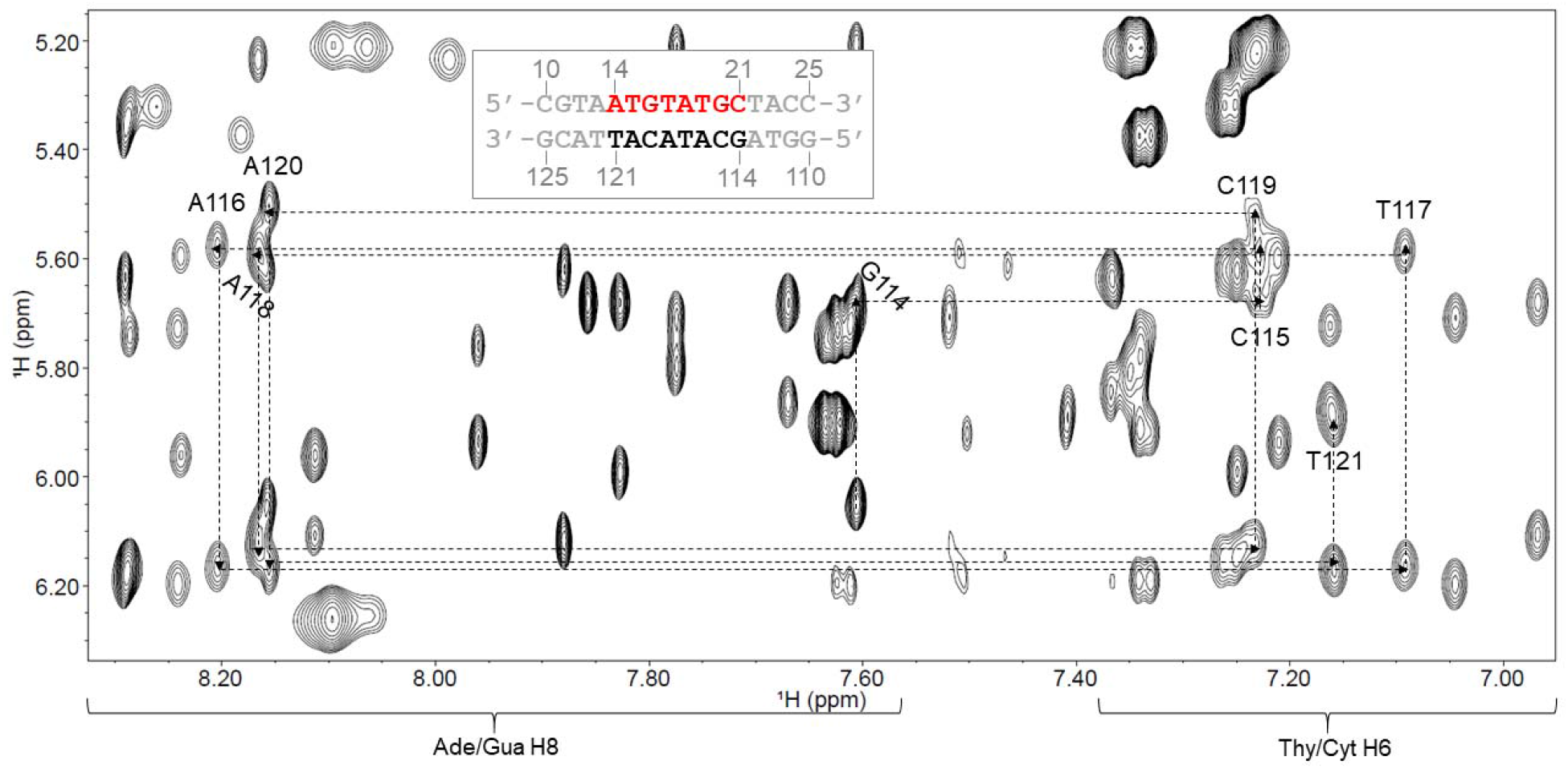
Base H6/H8 to H1’ region of the 2D NOESY spectrum illustrating sequential resonance assignments of the bottom stand (BS) of the 16-mer *loxP* spacer construct. Labels indicate the residue to which the H6 or H8 belongs. The numbering scheme is shown in the inset.

**Figure S2.**
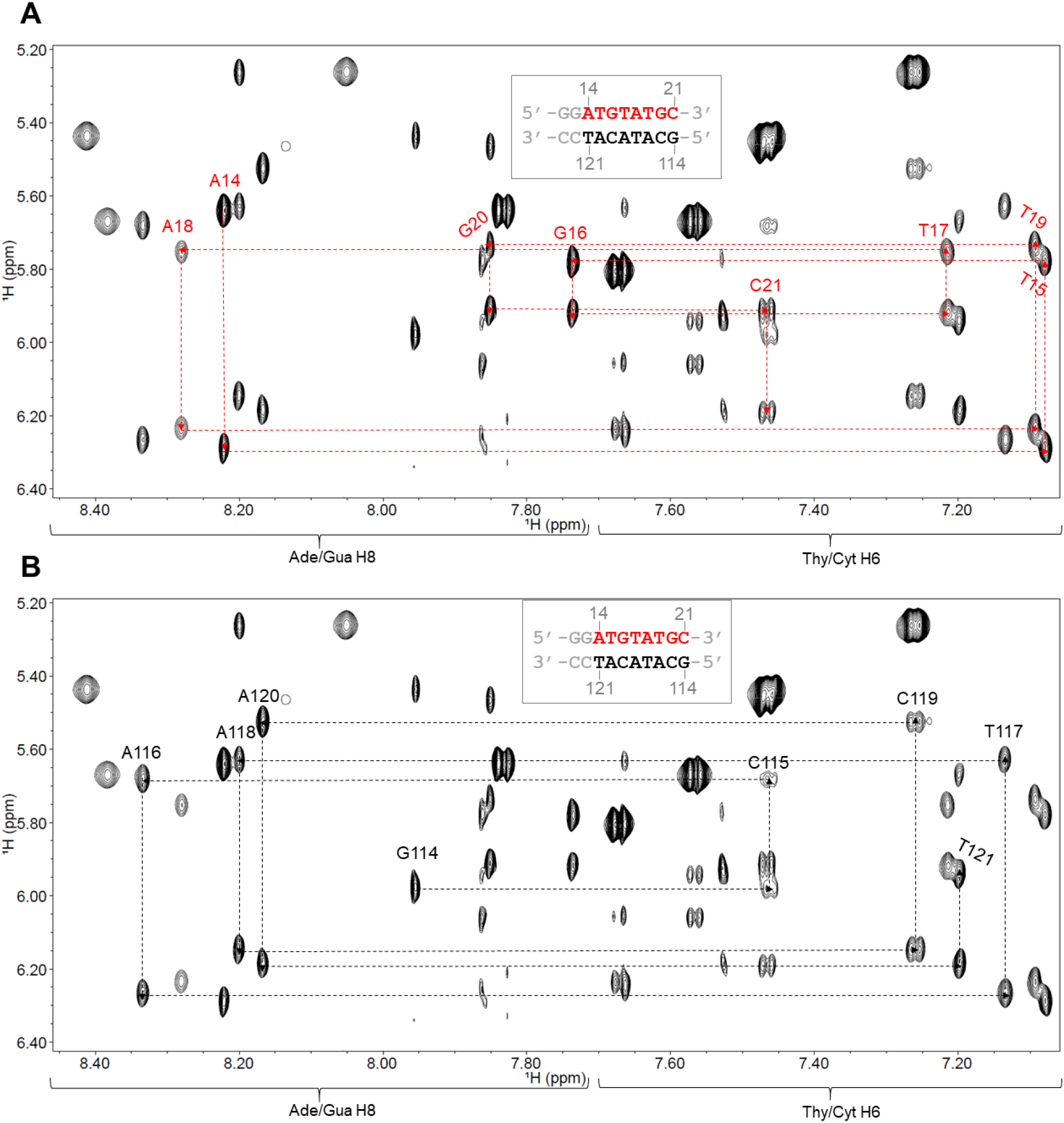
Base H6/H8 to H1’ region of the 2D NOESY spectrum illustrating sequential resonance assignments of the 10-mer *loxP* spacer construct. Labels indicate the residue to which the H6 or H8 belongs. The numbering scheme is shown in the inset. A) Top strand assignments. B) Bottom strand assignments.

**Figure S3.**
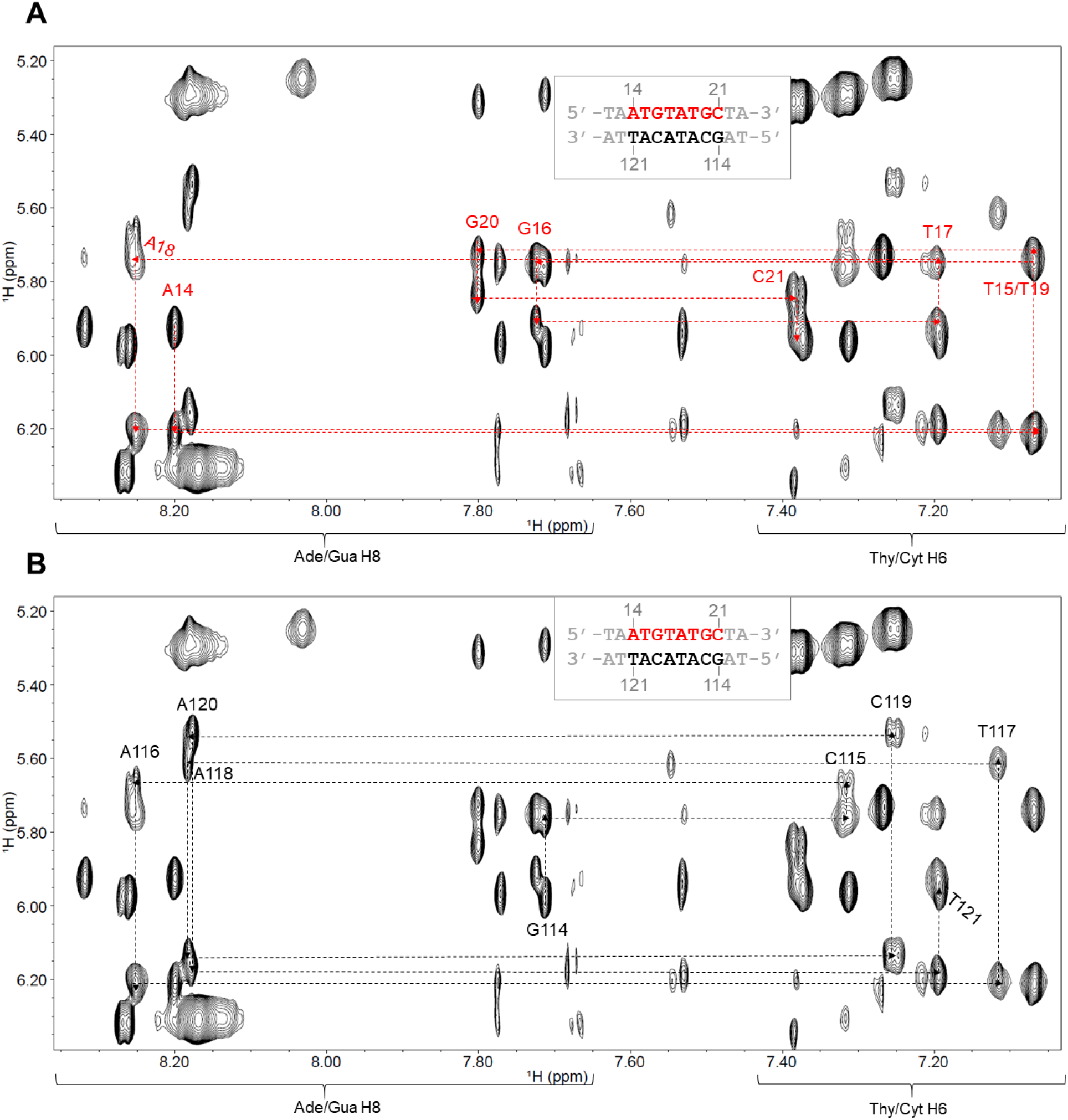
Sequential resonance assignments of the 12-mer *loxP* construct. Residue labels indicate the residue to which the H6 or H8 belongs. The numbering scheme is shown in the inset. A) Top strand assignments. B) Bottom strand assignments.

**Figure S4.**
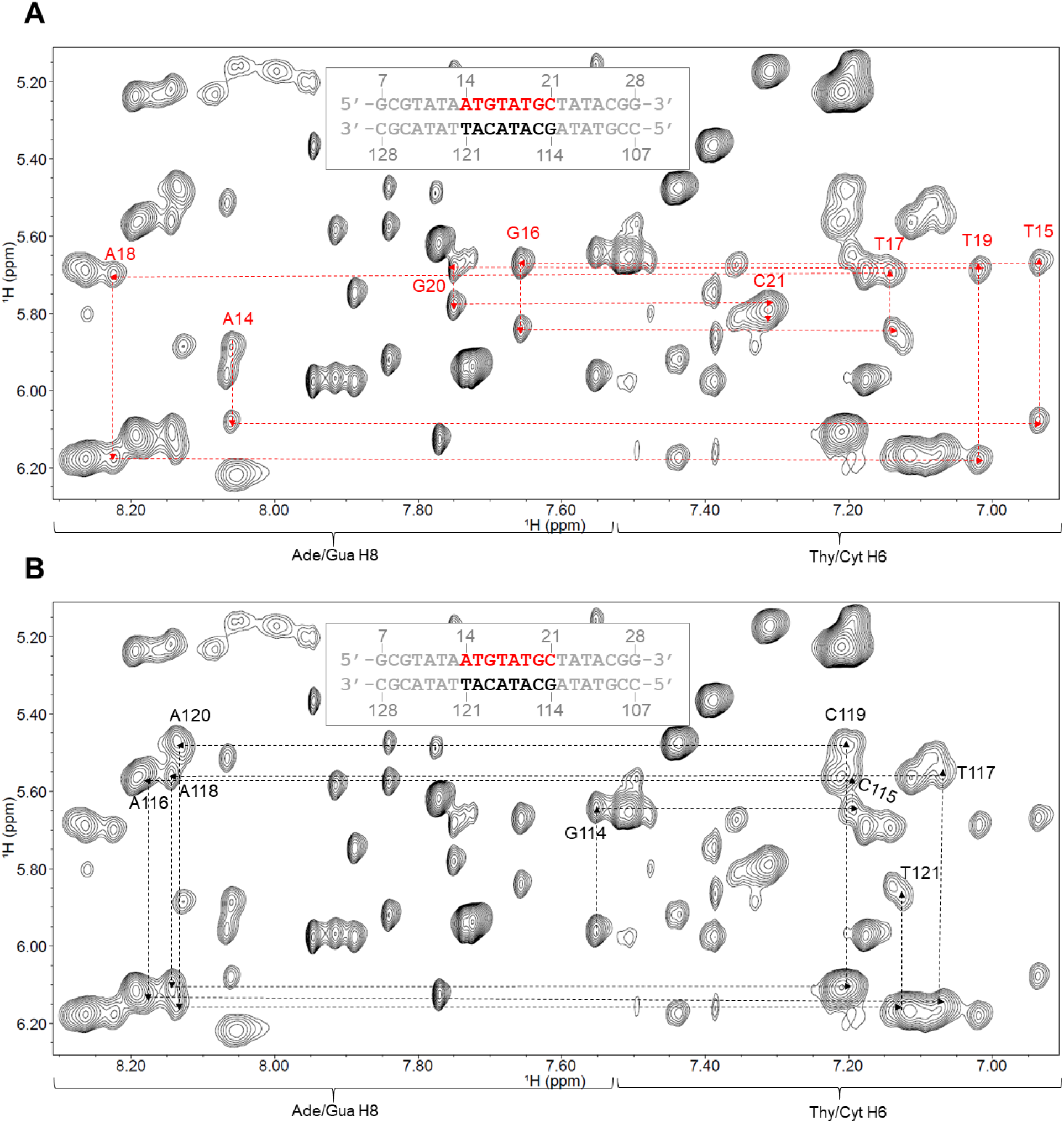
Sequential resonance assignments of the 22-mer *loxP* construct. Residue labels indicate the residue to which the H6 or H8 belongs. The numbering scheme is shown in the inset. A) Top strand assignments. B) Bottom strand assignments.

**Figure S5.**
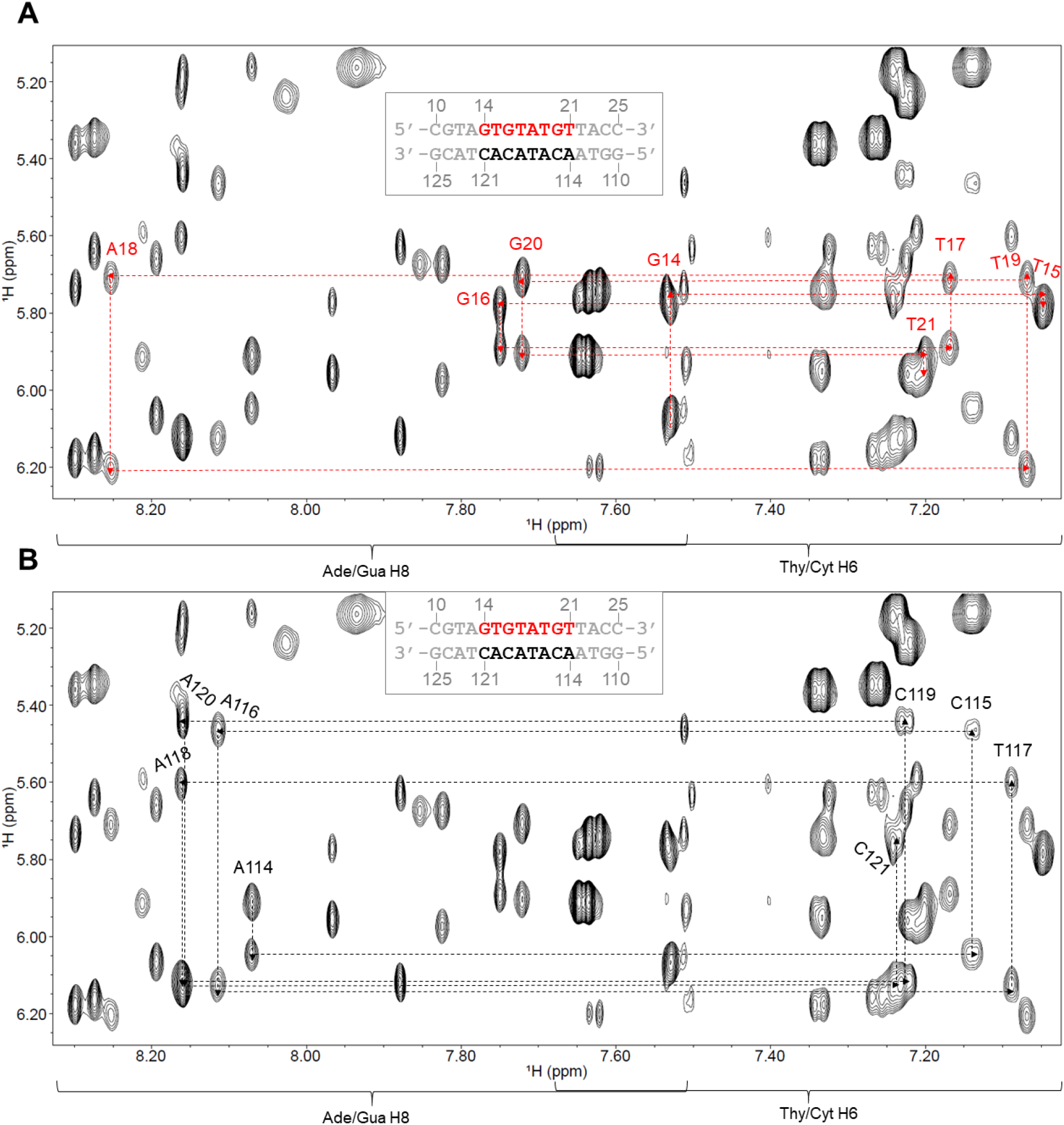
Sequential resonance assignments of the 16-mer *lox4* construct. Residue labels indicate the residue to which the H6 or H8 belongs. The numbering scheme is shown in the inset. A) Top strand assignments. B) Bottom strand assignments.

**Figure S6.**
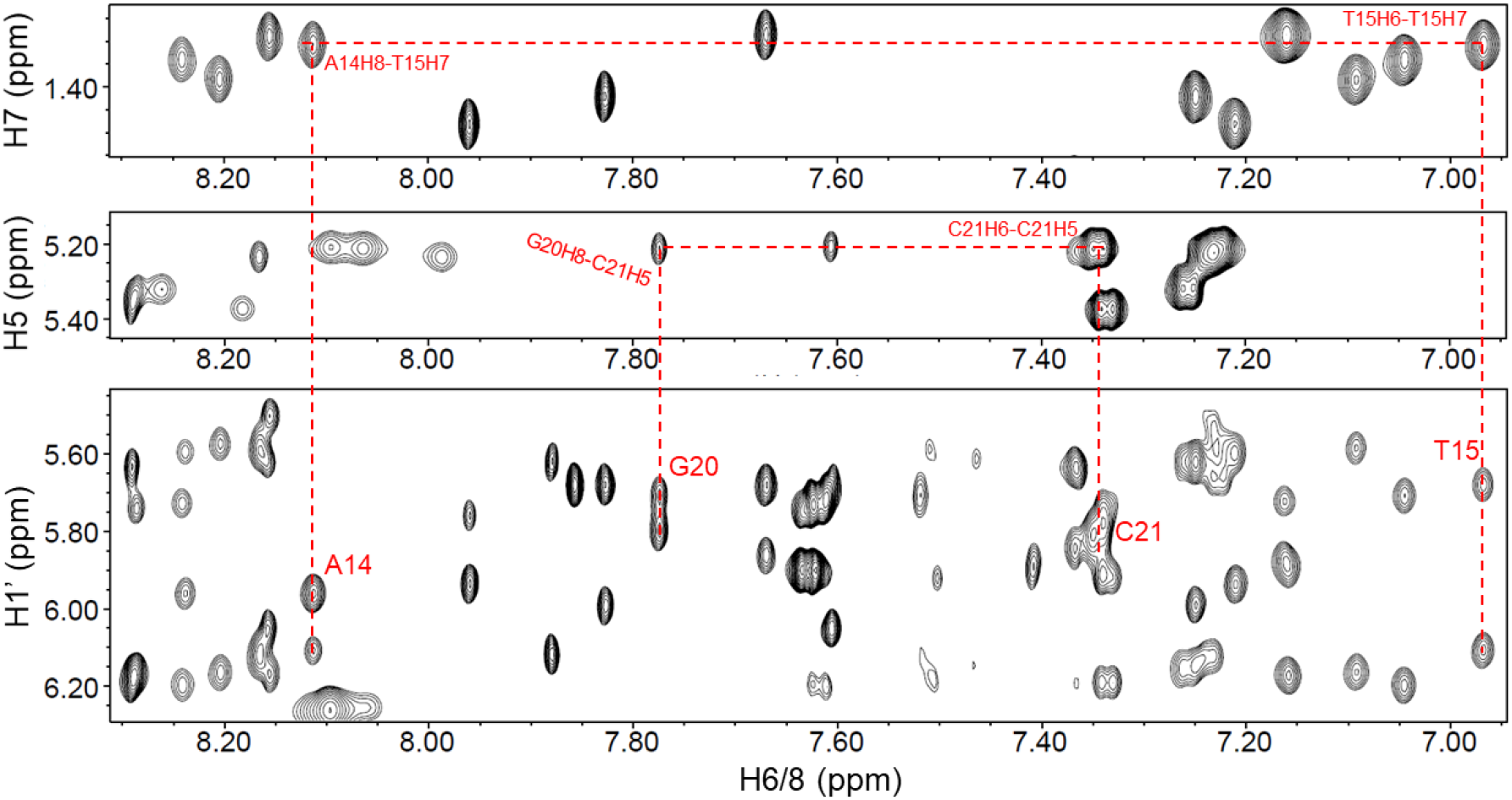
Use of thymine H7 and cytosine H5 protons to support sequential assignments. Each thymine H7 (methyl group) NOEs to its own H6 and to the H6/H8 of the preceding (5’) residue. Shown here, NOEs from T15H7 to both T15H6 and A14H8. Each cytosine H5 proton NOEs to its own H6 proton (with splitting due to J-coupling in the H6-H5 crosspeak) and to the H6/H8 of the preceding (5’) residue. Shown here, NOEs from C21H6 to C21H5 and G20H8.

**Figure S7.**
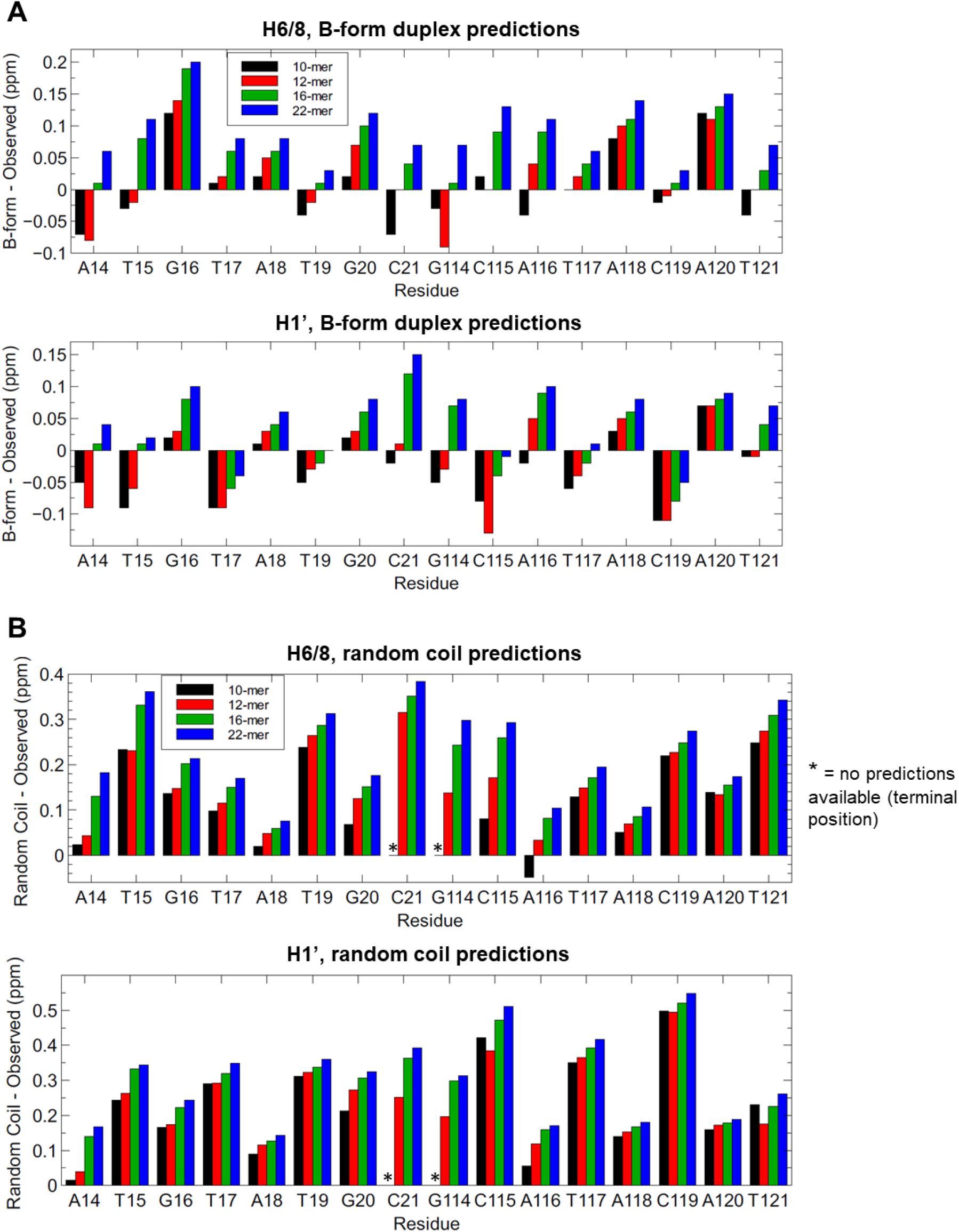
A) Differences between observed *loxP* chemical shifts and predicted double-helical B-form chemical shifts calculated with DSHIFT using the Altona method^26,27^. B) Differences between observed *loxP* chemical shifts and predicted random coil chemical shifts calculated with DSHIFT^26,29^.

**Figure S8.**
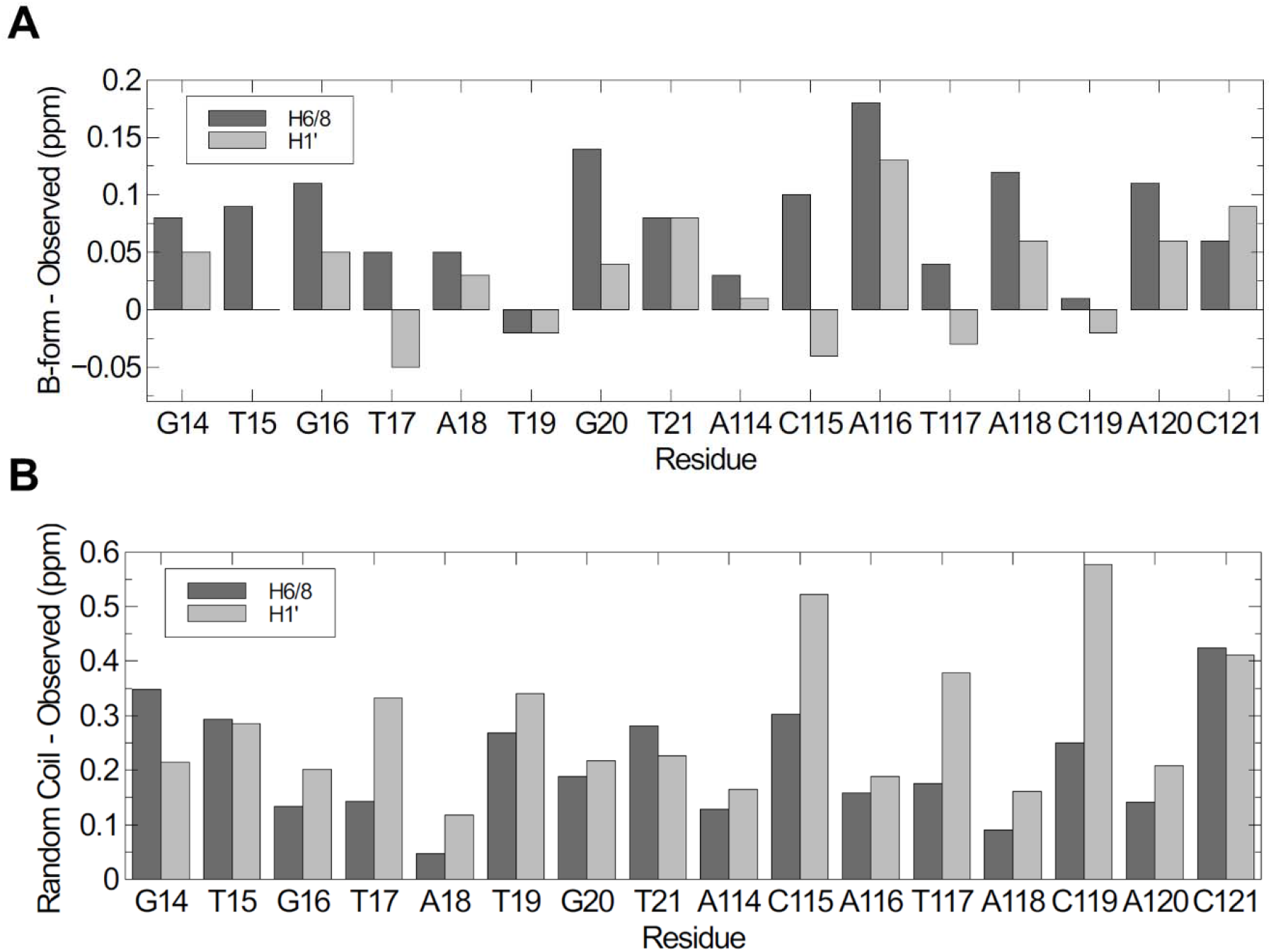
A) Differences between observed *lox4* chemical shifts and predicted double-helical B-form chemical shifts calculated with DSHIFT using the Altona method^26,27^. B) Differences between observed *lox4* chemical shifts and predicted random coil chemical shifts calculated with DSHIFT^26,29^.

## Notes

### Competing Interest Statement

The authors have declared no competing interest.

